# Disorder with consequence: Phosphorylation sites in HSPB5 yield distinct structural outcomes

**DOI:** 10.1101/2025.10.27.684587

**Authors:** Natalie L. Stone, Maria K. Janowska, Lucas Narisawa, Lisa M. Tuttle, Lindsey D. Ulmer, Miklos Guttman, Matthew F. Bush, Rachel E. Klevit

## Abstract

HSPB5, a member of the small heat shock protein family, acts as a first responder to cellular stress. One proposed mechanism of stress activation is phosphorylation. HSPB5 is phosphorylated at three sites—serine residues at positions 19, 45, and 59—located within its disordered N-terminal region (NTR). The extent of phosphorylation of the different sites leads to different cellular outcomes. HSPB5 forms polydisperse oligomers, where the NTR regions can either be exposed to the solvent or buried within the oligomer, forming internal contacts. We assessed the effect of single and triple phospho-mimicry on HSPB5 oligomeric properties. Our findings indicate that single phosphorylation causes localized and subtle changes in oligomer size, subunit exchange, hydrogen-deuterium protection patterns, and ability to delay aggregation of a known eye lens client, γD-crystallin. In contrast, the triple phosphomimic shows substantial structural and functional alterations. We provide a rationale for the increased chaperone activity observed in the S45D phosphomimic. Taken together, our results offer structural insights into how different phosphorylation events lead to distinct cellular outcomes.

## INTRODUCTION

Protein phosphorylation serves as a molecular switch that regulates protein activity, localization, stability, and interactions—allowing cells to rapidly respond to internal and external cues. As a reversible post-translational modification, phosphorylation modulates key processes such as signal transduction, cell cycle progression, and stress adaptation. Among stress-responsive proteins, the small heat shock protein HSPB5 (also known as αB-crystallin) is constitutively and ubiquitously expressed across tissues. It can be activated by environmental stressors or chemical modifications, including phosphorylation (reviewed in (1)).

HSPB5 contains three canonical phosphorylation sites—Ser19, Ser45, and Ser59—all located within its disordered N-terminal region (NTR). The kinases responsible for phosphorylating Ser45 and Ser59 are known, but the kinase that targets Ser19 remains unidentified (2, 3). The cellular functions or localization of phosphorylated HSPB5 is not uniform (2–8). Phosphorylated HSPB5 (pHSPB5) at position Ser19 and Ser45 localize to axons and dendrites with a filamentous-like staining pattern, whereas pSer59 is found in dendrites, especially along the plasma membrane and in spines (9). Disease or stress events also lead to specific phosphorylation patterns. In an animal model of stroke, phosphorylation of HSPB5 at Ser19 and Ser59 was present in neurons within the infarct area, but no modification at Ser45 was detected (4). Similarly, pS59 is enriched in astrocytes within active demyelinating multiple sclerosis lesions, whereas pS45 is constitutively present in astrocytes from normal-appearing white matter and remains unchanged in diseased regions (7). Thus, phosphorylation site usage is context-dependent and dynamically regulated by the cellular environment.

Phosphorylation is thought to induce conformational changes within HSPB5 oligomers (1). Historically, many studies have relied on a triple phosphomimetic variant (termed here “D3”) in which all three serines are mutated to aspartic acid or glutamic acid (10–12). Although D3 is a useful biochemical tool, it does not reflect physiological reality: simultaneous phosphorylation at all three positions is unlikely to occur in vivo. Moreover, the D3 model masks the nuanced—and often opposing—effects of individual or paired phosphorylation events. For example, in the context of chaperone activity during the biogenesis of multi-pass transmembrane proteins in cellular context, the phosphomimics S19D and S45D exert a strong inhibitory influence, both independently and in combination (8). In contrast, S59D does not inhibit and instead counteracts the suppression caused by phosphomimetic substitutions at the other two positions (8). In a study by Bartelt-Kirbach *et. al* double phosphorylation S45 and S59 protected dendritic trees complexity (13). Kuipers *et al*. (7) showed that site-specific phosphorylation alters HSPB5’s client interaction profile. Taken together, these results suggest that phosphorylation at different sites leads to distinct functional and structural outcomes (11, 12, 14, 15).

The biological findings highlight the need to define the changes within oligomeric ensemble caused by single phosphorylation and focus on site-specific consequences of HSPB5 phosphorylation. It is clear that even single-site phosphorylation can reshape oligomeric architecture in ways that redirect cellular targeting (4, 7)—but the structural and mechanistic underpinnings of these changes are still not well understood. In this study, we characterized the effects of single and triple phosphomimics (D1s and D3, substitution of serine to aspartic acid) on HSPB5 oligomers, when are polydisperse and highly dynamic. Our findings reveal that D1s induce specific and distinct changes within the oligomeric ensemble, while D3 produces a markedly different profile from WT. Among the single mutants, S45D showed the most pronounced perturbation, with increased exposure of the Long Hydrophobic subregion located in the N-terminal region (NTR) and enhanced chaperone activity against γD-crystallin aggregation. These results provide a foundation for defining how individual phosphorylation events modulate HSPB5 function through conformational remodeling.

## RESULTS

### Activation of HSPB5 chaperone activity by phosphorylation is site-specific

HSPB5 assembles into polydisperse oligomers stabilized by a quasi-ordered network of intra-oligomeric, inter-subunit interactions among its structured and disordered regions. These interactions involve the N-terminal region (NTR), α-crystallin domain (ACD), and C-terminal region (CTR), forming a complex interactome of NTR–NTR, ACD–NTR, and ACD–CTR contacts. Each site can engage multiple partners through diverse interactive patches, generating a highly interconnected architecture we refer to as quasi-order (16). This dynamic architecture enables transitions between inactive and active chaperone states in response to cellular signals (17–22). Among proposed modulators of this transition, phosphorylation is a particularly general and tractable mechanism. To investigate how phosphorylation influences HSPB5 function, we assessed the impact of phosphomimetic substitutions on its chaperone activity toward γD-crystallin. γD-crystallin is a lens protein prone to aggregation and is implicated in cataract pathology. HSPB5’s chaperone activity toward γD-crystallin depends on its NTR (23), which also harbors all three phosphorylation sites. Phosphorylation at each site has been observed in human lens (24).

Aggregation assays were performed in the absence of HSPB5 and in the presence of either WT, D1s (S19D, S45D, S59D) and D3 variants. We used the γD-crystallin-W130E mutant, which aggregates spontaneously at 37°C, to evaluate chaperone activity under physiological temperature and pH conditions (18, 23, 25). At the upper end of the lens-relevant range (∼pH 6.5–7.5; (26–28)), wild-type HSPB5 shows limited activity, so we used pH 7.5 to test whether phosphomimicry activates HSPB5 chaperone function. As shown in Fig. 1A, HSPB5-D3 robustly suppressed γD-crystallin aggregation over the 3-hour time course, whereas wild-type HSPB5 was ineffective. The S19D and S59D mutants behaved similarly to wild-type, whereas S45D showed enhanced activity. These results indicate that phosphomimicry effects are site-specific even within a disordered region and that S45 site phosphorylation (or mimicry) specifically increases chaperone function towards the γD-crystallin. The finding underscores that even within disordered regions, functional outcomes depend on precise molecular context.

**Figure 1.**
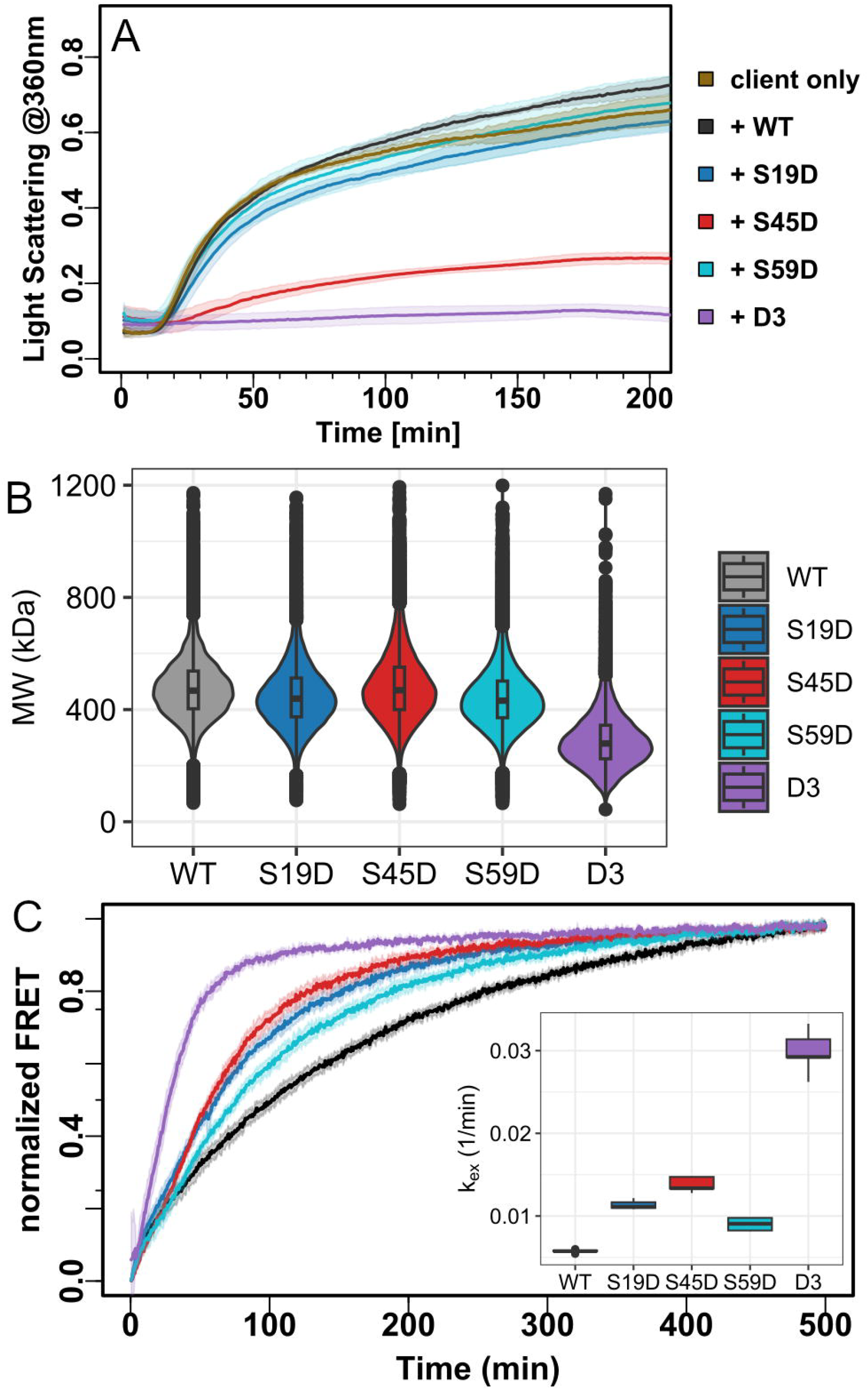
Comparison of wild-type HSPB5 and phosphorylation mimics (WT, S19D, S45D, S59D, D3). A. Chaperone activity measured by suppression of γD-crystallin aggregation. B. Oligomer size distributions from mass photometry. C. Subunit exchange kinetics assessed by FRET; inset shows boxplot of fitted k□□ values from five replicates.

### Macroscopic changes within the oligomers caused by the phosphorylation

A prevailing model for HSPB5 function is that its chaperone activity correlates with the size of oligomers present and/or the rate of subunit exchange between oligomers. We assessed the oligomeric sizes of HSPB5 and its phosphomimics by mass photometry (Fig. 1B, S1, Table S1). Consistent with polydisperse oligomers, HSPB5 and its variants exist in broad size distributions. All three D1s closely resembled WT in average oligomer size (WT, 477 ± 123 kDa; S19D, 453 ± 132 kDa; S45D, 487 ± 208 kDa; S59D, 446 ± 123 kDa) but D3-HSPB5 was markedly smaller (293 ± 109 kDa), similar to previously reported results (11, 12, 14, 15).

HSPB5 oligomers are dynamic assemblies in which individual subunits can dissociate and incorporate into other oligomers. Subunit exchange is typically measured using either FRET or fluorescence quenching assays, in which oligomers are prepared with fluorescently labeled subunits (29, 30). To streamline the workflow and avoid generating multiple labeled phosphomimetic mutants, we modified the standard protocol. For each sample, fluorescently labeled HSPB5 was mixed with unlabeled species (WT, D1s, or D3) at a final ratio of 1:5. This dilution strategy ensures that an average 24-mer contains ∼4 Alexa-labeled subunits and ∼20 unlabeled subunits. Subunit exchange was assessed by monitoring changes in the FRET signal upon mixing of donor- and acceptor-containing oligomer populations (Fig. 1C, S2, Table S1), and kinetics were fit to a first-order equation.

WT exhibited the slowest exchange rate, while D3 exchanged ∼5-fold faster. Among the single phosphomimics, S45D showed the highest rate (∼2.5-fold faster than WT), with S19D and S59D displaying intermediate behavior. Although overall exchange rates (k□□) were slower than in previous studies—likely due to differences in mixing strategy and assay conditions—the increase in subunit exchange rates for phosphomimics compared to WT is consistent with prior findings (10, 14, 30).

Integrating data from chaperone assays, oligomer size, and subunit exchange, we draw several conclusions: 1. Phosphomimicry in the NTR can alter oligomer organization. The D3 mutant shows significant perturbation in both size and exchange rate, consistent with disruption of the self-interaction network. 2. Although D1s do not display dramatic changes in oligomer size, they exhibit detectable increases in exchange rate. 3. Phosphorylation sites yield distinct functional outcomes—e.g., S45D enhances chaperone activity toward γD-crystallin, while neither of the other D1 mutants had a significant effect. 4. Chaperone activity cannot be predicted solely from oligomer size or subunit exchange. In sum, our data and literature findings suggest that size and subunit exchange should be interpreted as emergent properties of oligomeric ensembles, rather than direct indicators of chaperone activity. Finer-grained structural information is needed to fully resolve the changes induced by phosphorylation.

### Changes in sHSP exposure caused by phosphomimicry

Although all three phosphorylation sites reside within the disordered NTR, only S45D enhances HSPB5’s ability to delay γD-crystallin aggregation—highlighting the importance of positional context. This suggests that each modification drives distinct rearrangements within oligomers. Hydrogen-Deuterium Exchange Mass Spectrometry (HDX-MS), which can capture the dynamics of large, polydisperse assemblies, shows that the NTR is partially protected, the ACD is highly protected, and the CTR is largely exposed (18, 23, 29). In earlier work (18), we defined five NTR subregions—Distal, Aromatic, Conserved/Critical, Long Hydrophobic, and Proximal. Each phosphorylation site is in a different subregion: Aromatic (Ser19), Long Hydrophobic (Ser45), and Proximal (Ser59). For the ACD, regions flanking its surface grooves are of particular interest, as they mediate NTR contacts (Fig. S3).

To assess how each phosphomimic alters structure, we performed HDX-MS on WT, single phosphomimics (D1s), and triple phosphomimic (D3). We sampled HDX-MS at four deuterium exposure timepoints spanning 4 seconds to 20 hours, enabling detection of changes in both rapid and long-term protection. As a first step, isotope distributions were fitted to a unimodal binomial model (see Experimental Procedures). Overlapping peptides enabled us to reconstruct uptake profiles at single-residue or small-fragment resolution (Fig. 2A). This near residue-level fractional D-uptake (*f*D_avg_) is a proxy for the average exchange for all underlying conformational states of each region. All variants showed similar uptake patterns at the domain level: the CTR was fully deuterated within seconds and the NTR exchanged more slowly. Across all variants, the order of exchange for NTR subregions was Conserved/Critical < Aromatic ≈ Distal < Long Hydrophobic ≈ Proximal. The ACD was strongly protected at early times, but areas of deprotection emerged later, especially near the dimer interface groove. These results align with earlier studies (18, 23, 31) but provide finer spatial resolution of protection patterns.

**Figure 2.**
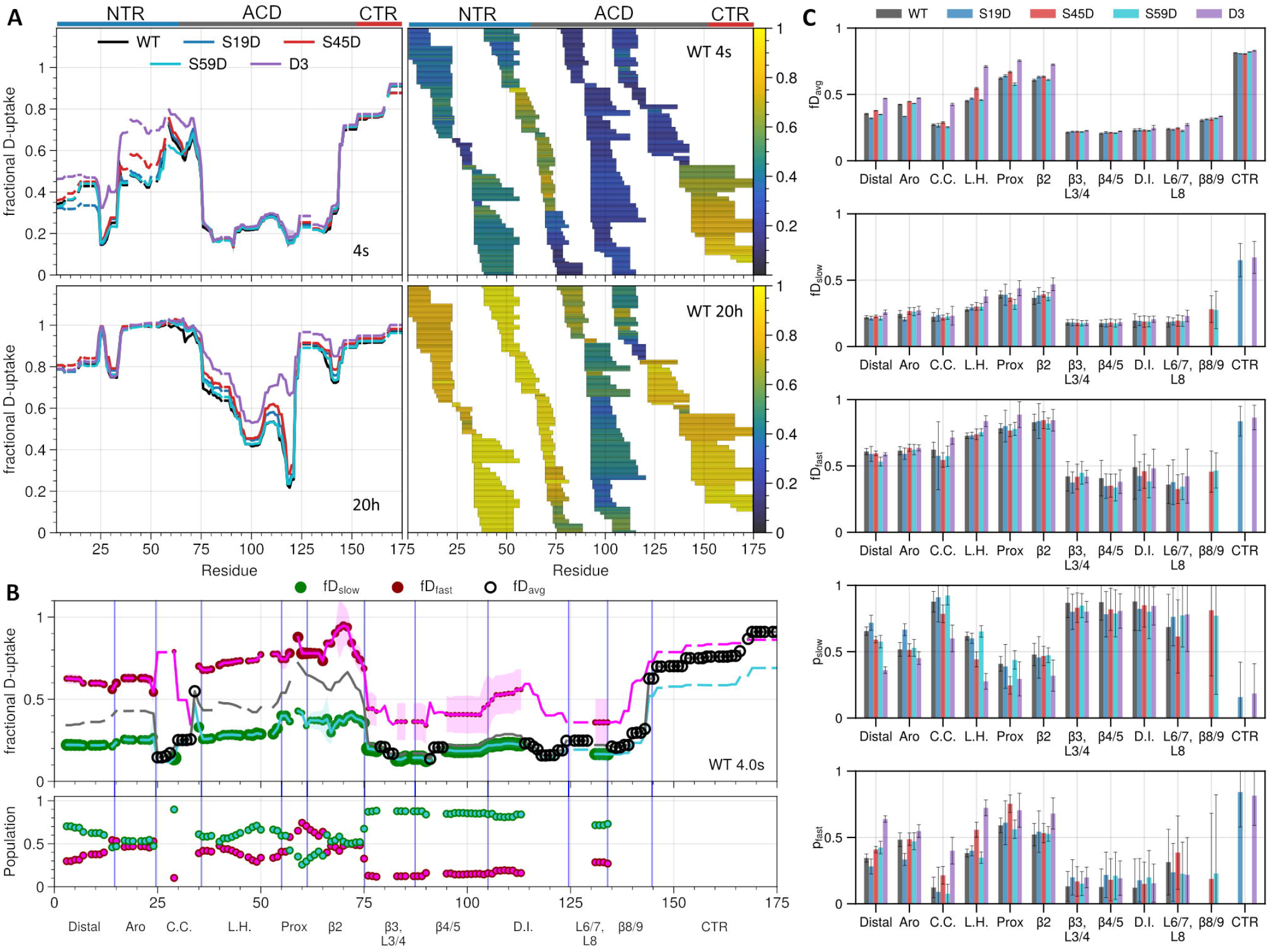
HDX-MS and bimodal behavior. A. Left, the per residue average fractional D-uptake (*f*D_avg_) for all variants based on unimodal fits of HDX spectra, shown for 4s top and 20h bottom. Right, peptide coverage maps for WT-HSPB5 with *f*D_avg_ colored using a blue to yellow gradient, shown for 4s top and 20h bottom. B. Unimodal or bimodal behavior of WT-HSPB5 at 4s. Top, lines show the 1- or 2-state fits of the data; circles indicate whether the unimodal (*f*D_avg_, black) or bimodal (*f*D_fast_, magenta and *f*D_slow_, green) fits are justified per residue. Bottom, corresponding populations (*p*_fast_, magenta and *p*_slow_, green) for residues with bimodal behavior. C. Bar plots showing all parameters per region defined in B, for the 4s exposure time. *f*D_avg_ values and error bars are calculated using all residues, whereas bimodal parameters only include residues where that fit is justified.

### Phosphomimic-induced region-specific changes

Each D1 species produced distinct local effects, whereas D3 caused widespread perturbations with nearly all NTR subregions deprotected early and ACD grooves exposed much earlier than in single mimics or WT (Fig. 2C and S5). S59D resembled WT the closest but showed unexpected protection near the mutation site. S19D stabilized the Aromatic region where it resides and at long exchange times it displayed increased exposure of ACD grooves, consistent with altered NTR–ACD contacts. Among single mimics, S45D showed the most pronounced perturbations with rapid deprotection of the Long Hydrophobic region where the substitution resides, of the Distal and Conserved/Critical subregions, and eventually, in ACD grooves (at 20h), consistent with NTR-ACD reorganization.

Substituting serine with aspartic acid increases the intrinsic exchange rate of the substituted amide and its neighboring C-terminal residue by ∼2-fold (32, 33). Thus, HDX-MS results must be interpreted carefully to ensure observed effects are not due solely to altered intrinsic exchange. In S45D and D3, the increases in exchange exceed what is expected from a Ser→Asp substitution alone, and these mutations affect peptides across the protein, not just those containing the substitution. By contrast, S19D and S59D effects are more localized (except for S19D’s influence on the ACD interface), and in both cases exchange actually decreases—opposite to the expected increase from a Ser→Asp substitution.

Together, these findings show that each phosphomimic has distinct effects on the exposure of specific NTR segments and thus on the network of interactions within the oligomer. The S45D variant exhibits the broadest effects across the NTR, perhaps explaining its increased chaperone activity towards aggregating γD-crystallin. The link between increased NTR accessibility and chaperone activity has been noted under other HSPB5 activating conditions, such as pH and disease-associated mutations (18). Interestingly, D3 shows dramatic solvent-exposure, greater than the sum of the perturbations induced by the single phosphomimics, indicating that dual- and triple-phosphorylation events may have synergistic effects.

### Slow and fast exchanging populations within HSPB5 oligomers revealed by HDX-MS deuterium uptake patterns

The quasi-order model of HSPB5 invokes a dynamic network of intra-oligomer interactions, where regions fluctuate between bound/interacting and unbound states. HDX-MS can capture this heterogeneity: peptides that exchange at two distinct rates (“slow” and “fast”) reflect populations in more-protected versus more-exposed environments. Bimodal analysis of peptide spectra enables identification of regions that exist in multiple environments and estimates of their relative populations (*p*_fast_ and *p*_slow_). The HDX-MS spectra of HSPB5 reveal many peptides that are visibly multimodal, even by eye. One example is shown in Fig. S4 for the NTR Long Hydrophobic peptide spanning residues 34–54, where a unimodal model poorly reproduces the experimental spectrum, but the bimodal model effectively reproduces the experimental spectrum a “fast” and “slow” exchange state. We use pyHXExpress and the approach previously applied to HSPB5 WT and disease mutants (31) to determine the bimodality across all residues for WT, D1s, and D3. The bimodal analysis complements the unimodal average exchange analysis: the unimodal analysis is associated with lower uncertainties and allows for broad comparisons while the bimodal analysis captures both dynamic and static heterogeneity. Together, the unimodal and bimodal analyses offer complementary views of structural dynamics and conformational heterogeneity.

The average fractional deuterium uptake (*f*D_avg_), along with values for the slow (*f*D_slow_) and fast (*f*D_fast_) populations, are depicted as a dark grey line and green and magenta lines, respectively and are shown for WT-HSPB5 at the 4s exposure timepoint in Figs. 2B and S4C. Filled green and magenta circles denote residues for which the bimodal fit was justified, whereas black open circles are denoted for residues where the data do not support fitting to additional populations. Bimodal behavior is observed across all three domains, the NTR displaying subregion-specific populations of fast- and slow-exchanging states, the ACD predominantly protected (∼80% slow), and the CTR largely exposed. This shows that even at the fastest exposure timepoint there are populations of NTR that are more protected, and populations of ACD that are less protected in the polydisperse HSPB5 oligomers.

The bimodal analysis reveals that phosphomimetic substitutions altered the balance between protected and exposed states in a region-specific manner (Fig. 2C and S5). Both S45D and D3 increased the proportion of fast-exchanging populations within the NTR, without changing the intrinsic uptake behavior of each state—except in the Long Hydrophobic region of D3, which exhibited elevated uptake at 4 seconds. In other words, the protection level of the fast and slow populations remained generally stable, but a greater fraction of subunits adopts the more exposed conformation.

The Distal, Conserved/Critical, and Long Hydrophobic regions all exhibited a higher proportion of fast-exchanging peptides in D3 compared to WT in the 4 second timepoint. S45D also shifted populations toward the exposed state: at 4 seconds, the Long Hydrophobic and Proximal regions were more deprotected, and by 60 seconds, this shift extended to the Distal and Conserved/Critical regions. These findings indicate that phosphorylation at that site does not globally destabilize the oligomer, but instead redistributes the ensemble of between protected and exposed states in specific NTR segments.

### The intra-oligomeric NTR interaction network is unchanged in activated phosphomimics

The S45D phosphomimic differs from the other D1 variants and WT in both chaperone activity and HDX-MS behavior, displaying both enhanced chaperone activity and a higher population of fast-exchanging regions. We hypothesized that the differences might reflect changes in the intra-oligomeric interaction network, and tested this using photo-crosslinking mass spectrometry (XL-MS). Previously reported photo-XL-MS characterization of WT-HSPB5 revealed numerous self-interactions between subunits (18, 23). To assess perturbation in these contacts due to phosphomimicry, we incorporated the photo-reactive amino acid p-benzoyl-L-phenylalanine (BPA) at specific NTR positions: W9B (Distal), F38B (Long Hydrophobic), and F61B (Proximal). These sites collectively interact with all NTR subregions and the two ACD binding grooves.

We generated mixed oligomers containing BPA-HSPB5 and either WT or S45D subunits at a 1:5 ratio, similar to the subunit exchange setup. XL-MS revealed interaction patterns consistent with prior data on WT-HSPB5, with W9B and F38B crosslinking to multiple NTR subregions and ACD grooves and F61B crosslinking to Distal and Aromatic regions.

Remarkably, comparison of the WT and S45D interactomes revealed no significant differences in interaction network (Fig. 3). Replicates for samples containing each of the BPA-containing variant exhibited no significant variation between crosslinks identified for WT and S45D (Fig. S6). To confirm that the oligomeric interactome remains unchanged between WT and S45D, we verified that crosslinks occurred between BPA-containing and non-modified subunits in the mixture approach. Because BPA and WT/S45D differ in molecular weight for several peptides, we were able to detect crosslinks between BPA-labeled and unlabeled proteins (Fig. S6). Taken together with bimodal analysis of the HDX-MS data, these findings suggest that S45D phosphomimicry does not alter the identity of self-interactions, but rather shifts the frequency or duration of contacts. In other words, the interactome remains intact, but the balance of protected versus exposed populations is redistributed—highlighting accessibility as a key determinant of HSPB5 function.

**Figure 3.**
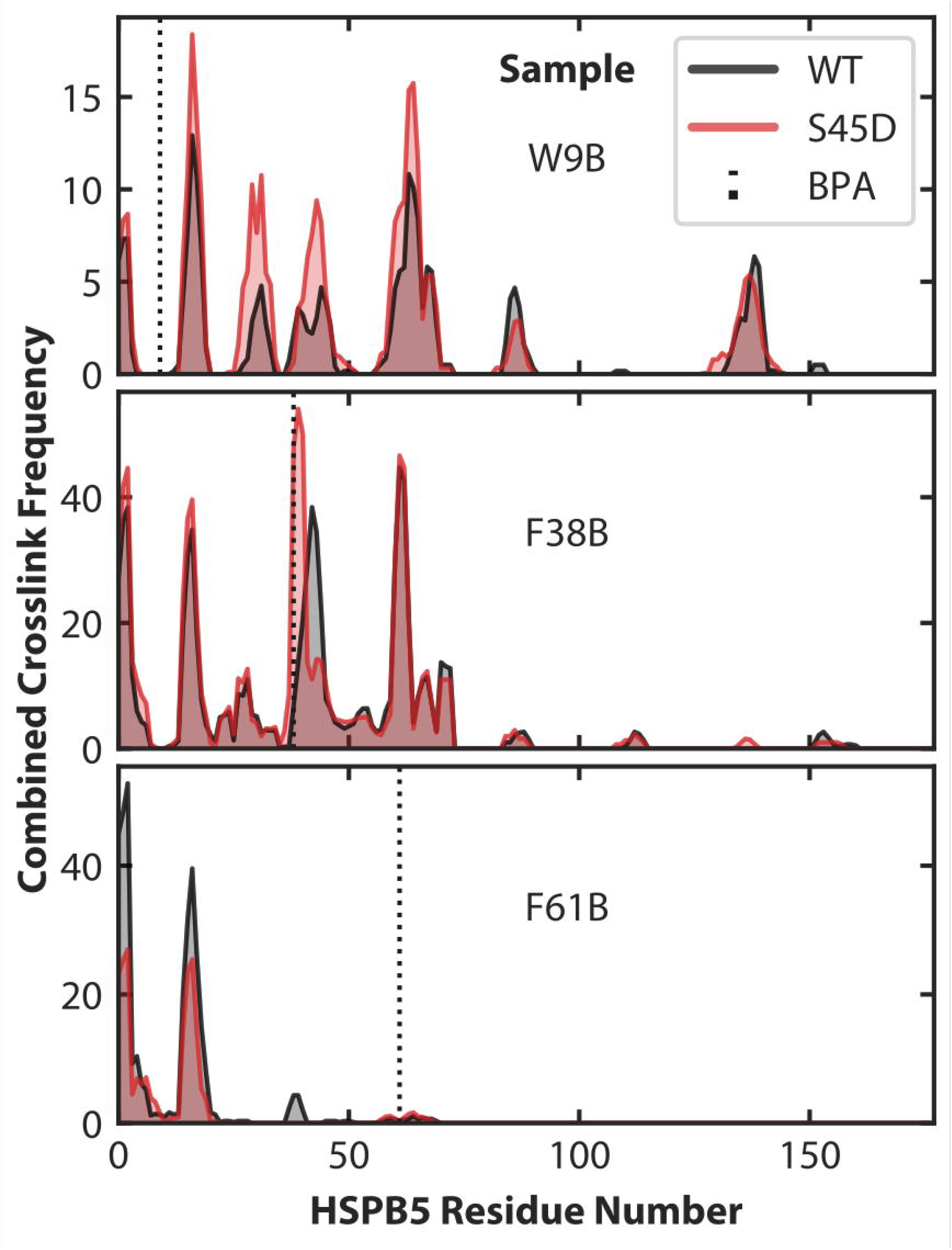
Counts of validated crosslinked spectra identified between BPA variants W9B, F38B, and F61B of HSPB5 to WT (black) or S45D (red) HSPB5. BPA variants (site indicated by dotted line) were individually diluted by WT or S45D in a 1:6 ratio, with three technical replicates each (Figure S6). A peptide spectral match (PSM) is the pairing between an experimental MS/MS spectrum and two covalently linked peptides. Results presented above sum the results from each replicate of a given sample, and a rolling average of PSMs across three residues are used to smooth crosslink distributions. For all samples, the WT and S45D crosslink identifications are correlated strongly.

**Figure 4.**
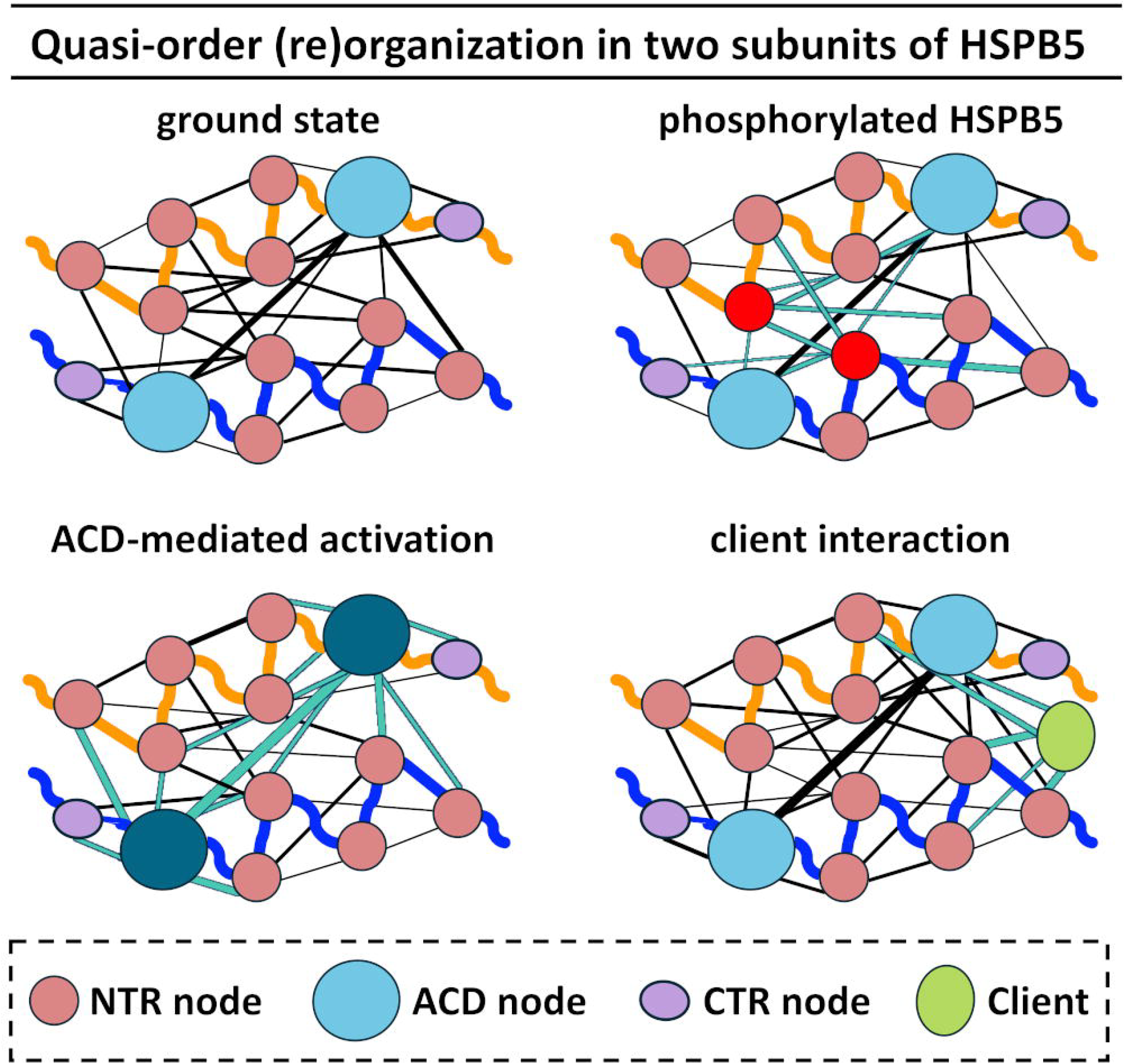
Subregions of the NTR and CTR, along with grooves on the ACD, have the capacity to interact both within sHSP dimers and across other subunits of the oligomer. Each interacting region can be conceptualized as a node within a broader interaction network. The strength of connections between nodes may be modulated by environmental stressors, mutations, and/or client binding. This figure illustrates potential changes in network connectivity between two monomeric subunits of HSPB5, while acknowledging that the full interaction network extends across additional oligomeric subunits. Backbone connectivity is represented by thicker curved lines in orange or blue. Interactions between nodes are depicted as straight lines. Modifications to the NTR (e.g., phosphorylation), the ACD (e.g., pH-dependent activation), or client binding to sHSPs can alter the strength of these node interactions. Network connections directly affected by such changes are highlighted in cyan; all other connections are shown in black.

## Discussion

HSPB5 is phosphorylated at Ser19, Ser45, and Ser59 within its disordered N-terminal region, with each site showing distinct regulation and localization in response to cellular stress conditions and across cell types. Simultaneous phosphorylation at all three sites is less common in cells, suggesting nuanced combinatorial, context-dependent regulation. Phosphorylated Ser19 and Ser45 localize to axons and dendrites, while phosphorylated Ser59 is enriched at dendrites (9) and dominates in Alexander’s disease pathology (34).

To assess structural consequences, we examined D1-HSPB5 variants. In HDX-MS experiments, S59D closely resembled WT, while S19D stabilized its local NTR and showed some ACD deprotection. S45D was the most perturbed, with increased exposure of the Long Hydrophobic NTR subregion and shifting the equilibrium toward solvent-accessible states in NTR and in ACD. Importantly, phosphorylation at Ser45 does not create new interaction sites but redistributes existing ones. At this point, the mechanistic basis for the distinct cellular outcomes and differing client preferences associated with phosphorylation at Ser19 and Ser59 remains unclear. What is clear is that each phosphomimic behaves as a distinct species, underscoring the need for site-specific structural analysis. Phosphomimicry likely underrepresents the full impact of true phosphorylation, thus to address these shortcomings further methodological development and experiments using fully phosphorylated proteins will be required.

The D3 triple mutant displayed the most dramatic changes, with widespread NTR deprotection and accelerated subunit exchange. We found the D3 oligomer to be smaller than WT. Conflicting conclusions among previous studies are likely due to concentration and solution conditions (11, 12, 14, 15). Our data support the prevailing view that D3 oligomers are destabilized and that subunit exchange is enhanced. D3 can serve as a model for probing high NTR/ACD exposure despite its unlikely existence in cells. But it is unlikely to provide insights into the more subtle and specific effects of the individual phosphorylated species.

Chaperone activity may or may not correlate with bulk oligomeric properties. Sometimes activity aligns with size, exchange, or hydrophobicity for a given client; other times, it does not. For example, we reported that all D1 variants showed similar bulk properties, yet only one has enhanced chaperone activity toward γD-crystallin.

Literature supports specificity of activation by client and site: in vitro studies of D1, D2, and D3 phosphomimics revealed client-specific outcomes—some had increased activity, others had decreased or left activity unchanged (11, 14, 35–37). These findings highlight that certain clients are protected by specific phosphorylation-induced activation modes.

Our findings demonstrate that phosphomimicry modulates HSPB5 function not by global destabilization or NTR untethering, but by rebalancing existing interactions, with direct consequences for oligomer architecture and client engagement. Bimodal HDX-MS analysis revealed that HSPB5 oligomers contain both protected and exposed regions, even in WT (31). At any time, approximately half of NTRs and ∼20% of ACDs populate more-exposed states, providing a reservoir for rapid client engagement.

Our data support a model of quasi-ordered oligomers as dynamic networks in which local perturbations can propagate globally: when a site becomes more or less interactive (as in S19D or S45D), the overall distribution of states shifts across the oligomer. The S45D interactome remains unchanged, but the populations of protected and exposed regions are altered. This behavior can be envisioned as a network of interconnected nodes each with different strengths of interaction. Changes in interactions between nodes can have far-reaching effects on other nodes leading to a reorganization of the entire network. Thus, even small changes in occupancy at individual sites can reverberate through the HSPB5 oligomers, reshaping the quasi-ordered landscape.

Finally, it is interesting to compare the alterations in the NTR as exemplified by the effects of phosphomimicry with alterations in the structured ACD as exemplified by disease-linked ACD mutations or pH-induced activation (18). Notably, both pH-induced activation and S45D phosphomimetic variant release the Long Hydrophobic NTR subregion and enhance chaperoning of γD-crystallin (18, 23). This convergence of outcome—despite distinct molecular origins—suggests that HSPB5 function tuned either by changes in the ACD or in the NTR reshape the quasi-ordered network and modulate the accessibility of client-binding surfaces. Together, these findings highlight two mechanistically distinct but functionally convergent modes of regulation, each capable of reorganizing the oligomeric interactome in response to physiologically relevant perturbations. As we study, sHSPs we expect to find more possible modes of action that stem from changes from the sHSPs quasi-ordered network.

### Experimental Procedures

#### Expression and Purification of HSPB5 WT and Mutants

Purification of HSPB5 variants, BPA-containing variants and γD-crystallin was performed as described previously (23). Briefly, *E. coli* BL21(DE3) cells were transformed with HSPB5 WT or mutants in a pET23 vector and expressed and purified using an untagged protein purification protocol including ammonium sulfate cut, anion exchange chromatography, and size exclusion chromatography previously (23). HSPB5-BPA-expressing cells were cultured identically to HSPB5 WT except for the addition of 1 mM *p*-benzoyl-L-phenylalanine (Bpa) shortly before induction, followed by induction with 1 mM IPTG and 0.02% (w/v) arabinose. HSPB5-BPA variants were purified similarly to WT, as previously described. γD-crystallin W130E variant was expressed in *E. coli* BL21(DE3) cells as a C-terminal HexaHis-Sumo fusion in pET28a vector and purified as previously described, by including immobilized metal ion affinity and size exclusion chromatography (23). The HexaHis-Sumo tag was removed by SENP1 protease cleavage followed by further purification. Concentrations of WT HSPB5, phosphomimic variants, and γD-crystallin W130E were determined by absorbance at 280 nm on a NanoDrop One spectrophotometer, while concentrations of BPA-containing HSPB5 were determined by BCA assay using a BSA standard curve.

### γD-crystallin chaperone activity assays

γD-crystallin chaperone activity assays were conducted as described previously (18, 23). Briefly, 100 uM of each HSPB5 variant was preincubated for >3 hours in 25 mM NaPi buffer, 150 mM NaCl, pH 7.5 at 37 °C. Following pre-incubation, 15 uL of each variant or buffer control was combined with 35 uL of buffer the wells of a Corning 96-well, half-area, flat-bottomed clear assay plate and briefly incubated again at 37 °C before addition of 50 uL of 600 uM ice-cold γD-crystallin W130E in 25 mM NaPi, 150 mM NaCl, 2 mM EDTA, pH 7.5 buffer. The plate was briefly mixed and then incubated at 37 °C in a BioTek Synergy HT plate reader where light scattering at 360 nm was recorded for several hours. Experiments were performed in triplicate. Final assay conditions in the plate were 300 uM γD-crystallin W130E and 15 uM HSPB5 variant in 25 mM NaPi, 150 mM NaCl, 1 mM EDTA, pH 7.5 buffer with a final well volume of 100 uL.

### Mass Photometry

Mass photometry experiments were conducted using the Refeyn TwoMP instrument. Data calibration was performed using beta-amylase (producing calibration peaks at 56, 112, and 224 kDa). Data acquisition was done using AcquireMP, and analysis was carried out with DiscoverMP. The experiments were performed using a buffer-free focusing method. The final concentration of HSPB5 variants was 1 µM in 25 mM NaPi, 150 mM NaCl, pH 7.5. Samples were incubated for 3 hours at 37°C before the experiment.

### Subunit exchange measured by fluorescence

Alexa Fluor 488 and Alexa Fluor 568 maleimide labeling protocol was described previously (29). 250 nM Alexa labeled S153C HSPB5 proteins were mixed with 1.25 μM unlabeled protein variants: WT, S19D, S45D, S59D, and D3, and incubated separately overnight at 37 °C to ensure full subunit exchange of the labelled variants with the unlabeled proteins. Samples of the donor and acceptor mixtures (Alexa488-unlabelled protein variant and Alexa568-unlabelled protein variant), were combined in 1:1 ratio just prior to the experiment in preheated plate. We observed the subunit exchange using FRET based assay with excitation and emission wavelengths of 488 nm and 568 nm, respectively. The FRET signal here is a proxy for the subunit exchange rather than actual distance measurement. Five replicate experiments were performed on a BMG Clariostar Plus plate reader. The subunit exchange rate was determined assuming first order reaction kinetics, fitting to the equation y = C1 – C2*exp(-*k*_1_t) (38).

#### Hydrogen-Deuterium Exchange Mass Spectrometry

The HDX-MS experiments were performed as previously described with some modifications listed below (18). In short, 20 uM samples of all HSPB5 variants were incubated at 37 °C for 3 hours in 25 mM NaPi, 150 mM NaCl, 1 mM EDTA, pH 7.5 buffer prepared in Optima MS-grade water and then cooled to room temperature prior to deuteration experiments. HSPB5 WT and phosphomimic variants were diluted 10-fold in in deuterium oxide-based PBS buffer (final samples: 2 uM protein in 85% D2O; pH* of D2O-based PBS buffer was 7.56) and incubated at room temperature for 4 seconds, 1 minute, 30 minutes, or 20 hours. At the end of the given timepoint, exchange was quenched by addition of an equal volume of ice-cold quench buffer (0.2% formic acid, 0.24% trifluoroacetic acid) to reduce sample pH to ∼2.5 and samples were immediately flash frozen in a dry ice/ethanol bath. Samples were stored at -80 °C until immediately before mass spectrometry analysis. Undeuterated control samples were prepared by the same method, except that protein stocks were diluted into H2O-based buffer, rather than D2O-based. Fully deuterated control samples were made by preparing 20 uM protein stocks in a 50:50 mixture of H2O-based PBS 7.5 and 7 M urea (final urea concentration of 3.5 M) and incubating at 90 °C to maximally unfold the protein, quenching, and storing as for the other samples. Samples were automatically thawed, digested using an immobilized Nepenthesin-2 protease column (AffiPro), peptides were collected on an XSelect CSH C18 trap cartridge (Waters) before injection onto an ACQUITY CSH C18 50 mm UPLC column (Waters), and then injected on a Waters Synapt G2-Si Q-TOF mass spectrometer using a setup built in-house around the LEAP PAL system (39) (Watson et al., 2021). Samples were analyzed in triplicate. Prior to HDX-MS dataset acquisition, HSPB5 peptides from undeuterated samples were identified by MS/MS on a Thermo Orbitrap Fusion Tribrid instrument and MS^E^ on a Waters Synapt G2-Si followed by data analysis using Byonic (Protein Metrics). Deuterium uptake was analyzed in HDExaminer 3.0 (Sierra Analytics), HDXBoxeR (40), and pyHXExpress (31). To allow the access to the HDX-MS data of this study, the HDX data summary table and the HDX data table are included as supporting information as per consensus guidelines (41).

The bimodal analysis follows the procedure described previously using pyHXExpress (31) and pyHDX (42) for near residue-level D-uptake calculations. Briefly, the average D-uptake (*f*D_avg_) is calculated based on unimodal fits of all peptides. Bimodal fits are also performed for all peptides where *f*D_fast_ is defined as the more exchanged state and *f*D_slow_ is the more protected state. For each residue, these data are aggregated over all peptides and charge states that contain that residue, the averages and associated errors are determined, and the populations *p*_fast_ and *p*_slow_ are calculated based on the relative distance of *f*D_fast_ and *f*D_slow_ to *f*D_avg_. To justify a bimodal fit the following criteria were applied: 1) *f*D_avg_ must fall between *f*D_fast_ and *f*D_slow_, 2) *f*D_fast_ and *f*D_slow_ must be non-overlapping including error bars, 3) the minimum allowed population must be greater than 10%, and 4) the errors on *f*D must be less than 0.25. Otherwise, the data does not support the bimodal fit and the unimodal fit is selected as appropriate for that region.

#### BPA Crosslinking Mass Spectrometry

The BPA crosslinking experiments were performed as described previously (23, 43) with some modifications. The samples contained 25 μM BPA HSPB5 mixed with 125 μM unlabeled protein variants WT- or S45D-HSPB5, and were incubated for 3h at 37 °C prior to UV crosslinking. The crosslinking reaction was performed for BPA sites 9, 38, and 61. Data was collected with an Easy Nano LC coupled to a Thermo Orbitrap Fusion Lumos Tribrid. Data was collected using data-dependent acquisition with a 30-minute gradient for samples containing HSPB5 W9B and F61B, or a 50-minute gradient for samples containing F38B (43). The mass spectrometry proteomics data have been deposited to the ProteomeXchange Consortium via the PRIDE partner repository (44) with the dataset identifier PXD069703.

#### Identification of crosslinks using the Trans Proteomic Pipeline (TPP)

Crosslinks were identified using the previously reported informatics workflow (43) utilizing the Trans Proteomic Pipeline version 7.2.0 (45). Briefly, the method uses Comet (46) to search for non-crosslinked peptides in each dimeric reactant sample and uses PeptideProphet (47) to assign false discovery rates (FDR) at the PSM level. In crosslinking, a peptide spectral match (PSM) is the pairing between an experimental MS/MS spectrum and two covalently linked peptides. Results are then used to filter the validated protein database to a 1% FDR with a minimum of two PSMs per protein, which is used as the protein database to analyze the dimeric product sample. Dimeric product samples are searched using Kojak (48) and the validated protein database. PeptideProphet is used again to assign FDR thresholds and filter results to a 1% FDR. For histograms, each PSM was associated with the residue that was assigned the highest probability of participating in a crosslink with BPA. When more than one residue was assigned the same probability, an equal fraction of that PSM was assigned to each of those residues. The code used to assign residue-level crosslinks and create histograms was modified from previous work (43) and is publicly available (https://github.com/bushgroup/Identifying-Site-Specific-Crosslinks).

## Supporting information

Supplemental_Table_HDXMS

## Abbreviations

sHSP: (small heat shock protein)
NTR: (N-terminal region)
ACD: (alpha-crystallin domain
CTR: (C-terminal region)
DLS: (Dynamic Light Scattering)
CD: (Circular Dichroism)
D1: (single phosphomimic)
D3: (triple phosphomimic)
EM: (electron microscopy)
HDX-MS: (Hydrogen-Deuterium Exchange Mass Spectrometry)
pHSPB5: (phosphorylated HSPB5)

## Acknowledgements

We would like to thank Christopher N. Woods for preforming preliminary HDX-MS experiments, Mia Cervantes for careful reading of the manuscript.

## Funding and additional information

Funding was provided by the National Eye Institute: 2 R01 EY017370 (R.E.K.) & T32 EY07031 (M.K.J.), National Institute on Aging: T32 AG066574 (M.K.J.), National Institute of General Medical Sciences: T32 GM153507 (L.N.), the National Institute on Aging through T32 AG066574 (L.D.U.), the University of Washington’s Proteomics Resource (UWPR95794). The content is solely the responsibility of the authors and does not necessarily represent the official views of the National Institutes of Health.

