## Supplementary_Info for "Disorder with consequence: Phosphorylation sites in HSPB5 yield distinct structural outcomes"

Materials include:

Extended Experimental Procedures

1 table

6 figures

**Extended Experimental Procedures**

**Extended LC-MS Methods**.

The following methods have been adapted from previous work (1). Data was acquired using an EASY NanoLC coupled to a Thermo Orbitrap Fusion Lumos Tribrid. Samples containing W9B or F61B were resuspended using 95% water and 5% ACN with 0.1% FA. Samples containing F38B were resuspended using 70% water and 30% ACN with 0.1% FA. The weight (μg) of protein loaded onto the gel and the relative color intensity of the band excised from the gel prior to digestion was used to estimate the weight of the digested peptides loaded on the column. For example, if 7.6 μg of protein was loaded onto a gel lane and the dimeric product band was roughly 30% of the total color intensity, we estimated that the excised dimeric products contained about 2.3 μg of protein. In that case, about 20% (500 ug) of the digested peptides would be loaded on the column. Roughly 500 ng of peptide was loaded onto an 8-cm trap column. The sample was then separated on a 25-cm analytical column with a 75 μm inner diameter using a 30-minute gradient for W9B or F61B samples, or 50 minutes gradient for F38B samples. The 30-minute gradient was from 6% B to 45% B, where A was water and B was 80% ACN, at 300 $nL \cdot min^{-1}$. The column was then flushed and regenerated for 35 minutes. The 50 minutes gradient was in two steps, a 25 minutes gradient from 5% B to 25% B, and another 25 minutes gradient from 25% B to 100% B, with the same solvent used as A and B. The column was then flushed and regenerated for 30 minutes. In the 30 minutes gradient, a charge state range of 2-8+ was utilized with 30 second dynamic exclusion, while the 50 minutes gradient utilized a 4-8+ charge state range with 10 seconds of dynamic exclusion. Spectra were acquired across the entire length of LC methods using data-dependent acquisition and an intensity threshold of $2.0 \times 10^4$. Orbitrap detection and higher-energy collisional dissociation (HCD) fragmentation (30% normalized collision energy) were used with a target value of $1.00 \times 10^5$, maximum injection time of 22 ms, top N of 20, and isolation width of 1.6. MS1 were acquired at a resolving power of 120,000 over the range of 400-2000 m/z, and MS2 were acquired with a resolution of 15,000. Mass spectra were acquired using profile mode and the Thermo RAW format. Those files were centroided and saved as mzml files using the ProteoWizard msconvert tool as implemented in the Trans Proteomic Pipeline version 7.2.0 (2).

**Comet Search Settings**. Details on Comet (3) search settings.

The settings used have been previously reported (1). The searches were enzyme nonspecific using a peptide mass tolerance of 20.0 ppm. The isotope error offset was 3, and BPA was defined as an additional amino acid, B, that has a mass of 251.09462859 Da. Methionine oxidation and cysteine iodoacetamide alkylation were variable modifications.

**Kojak Search Settings**. Details on Kojak (4) search settings.

The settings used have been previously reported (1). The search settings matched those described for the Comet searches except that the precursor tolerance was 15 ppm and enzyme selection rules were used. For all samples, the cleavage sites of D and E were added to the trypsin settings to match the dual-enzyme trypsin-GluC digestion. Crosslinks were defined as from BPA to any residue, with no change in mass.

**PeptideProphet Settings**. Details on PeptideProphet (5) settings.

The settings used have been previously reported (1). PeptideProphet was used to validate both Comet and Kojak results with the following options: ppm for accurate mass binning, only use Expect Score as the discriminant, decoys to pin down the negative distribution, use non-parametric model, and report decoy hits with computed probability. For all samples, the enzyme also was defined in the additional options line as -e"TrypGluC:specific:true:KRDE|P".

For Comet searches, after filtering using a 1% False Discovery Rate (FDR) and a minimum of 2 Peptide Spectral Matches (PSMs), this yields a protein database for the sample. A 1% FDR is defined as the error rate of 0.0100 that PeptideProphet reports in the error table. This is a PSM level FDR. PSMs that meet the 1% FDR threshold are found by filtering to spectra that have a PeptideProphet probability greater than or equal to that for an error rate of 0.0100. For Comet, overall (considering all charge states) error rates, FDRs, and PeptideProphet probabilities were used. For Kojak searches, the same PeptideProphet settings were used, but for ions of each charge state, the corresponding error rate, FDR, and PeptideProphet probability were used.

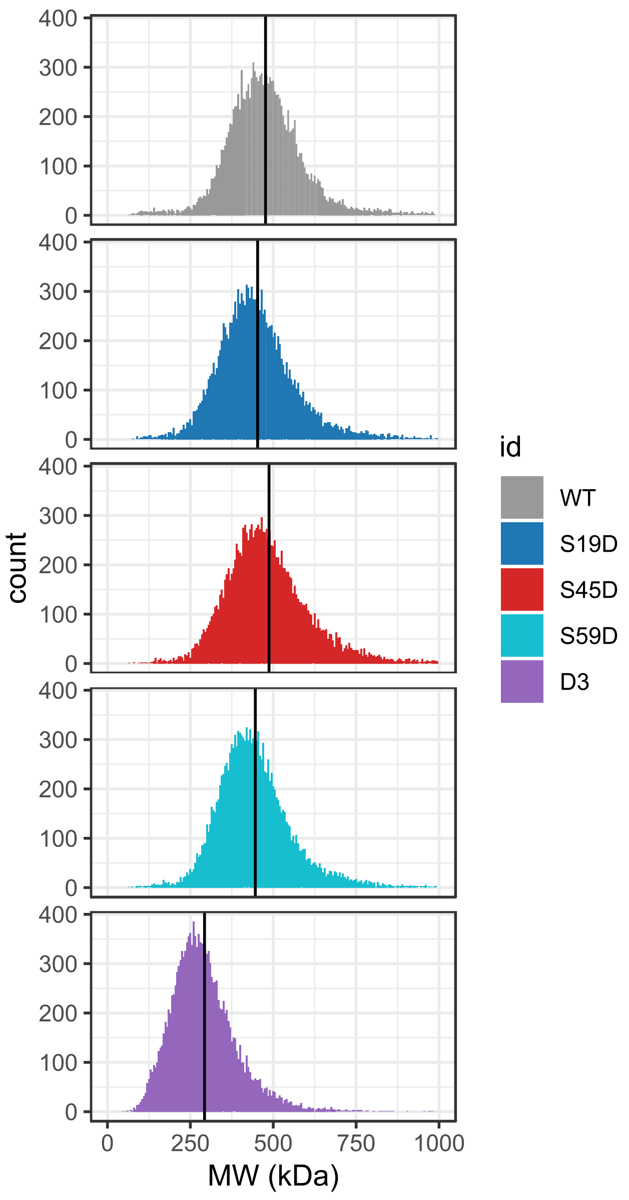


Figure S1. Mass photometry size distribution for different protein variants. The mean values for each distribution are shown as a black line.

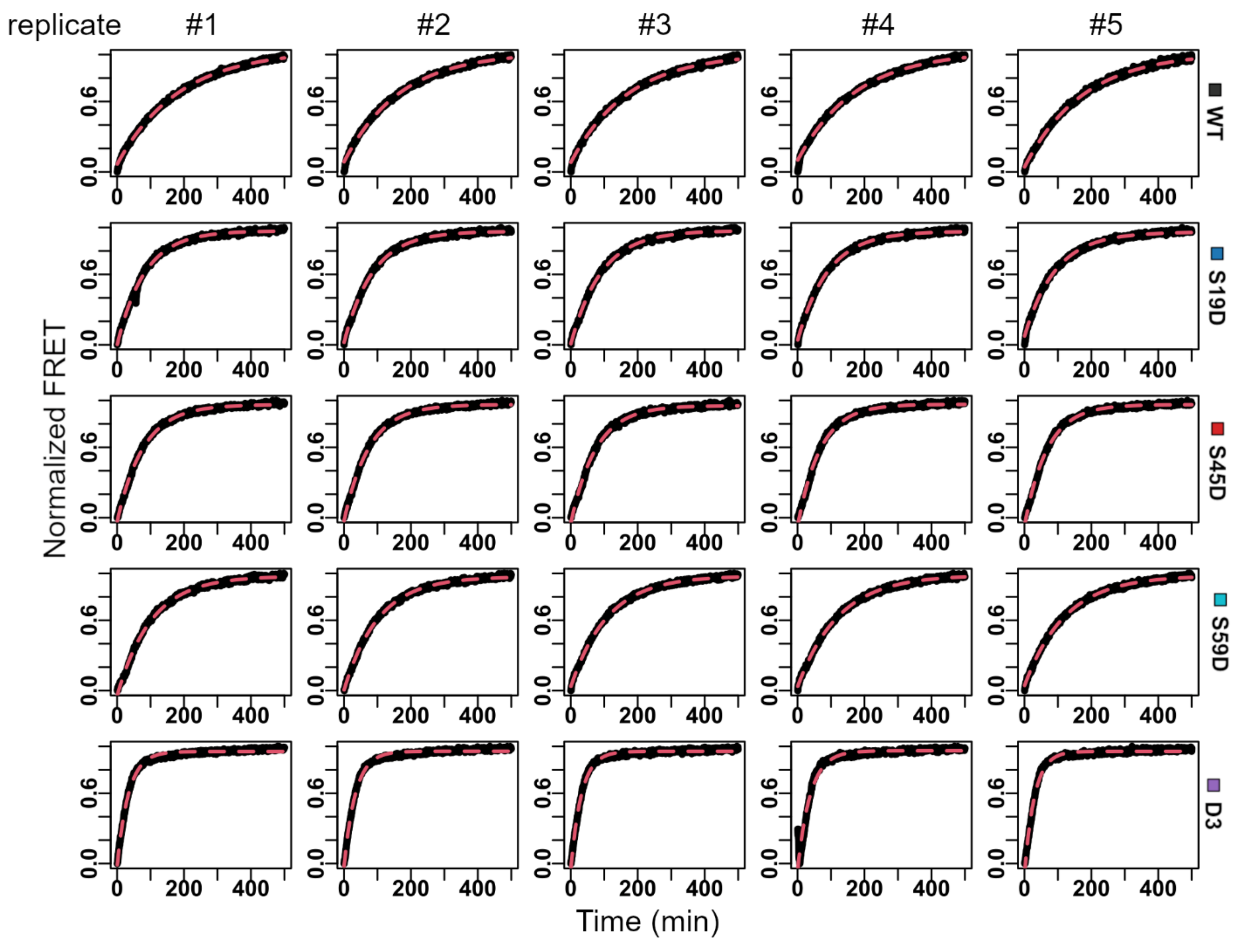


Figure S2. The subunit exchange fits for 5 replicates (columns) for each protein variants (rows) from FRET experiments.

| **Protein** | **MW (kDa)** | **$k_{ex}$ (1/min)** |
|---|---|---|
| **WT** | 477±123 | (5.7±0.20) x $10^{-3}$ |
| **S19D** | 453±132 | (11±0.05) x $10^{-3}$ |
| **S45D** | 487±208 | (14±0.08) x $10^{-3}$ |
| **S59D** | 446±123 | (9.0±0.07) x $10^{-3}$ |
| **D3** | 293±109 | (30±0.26) x $10^{-3}$ |

Table S1. Molecular weight determined by mass photometry and the subunit exchange rates determined by FRET for the HSPB5 variants.

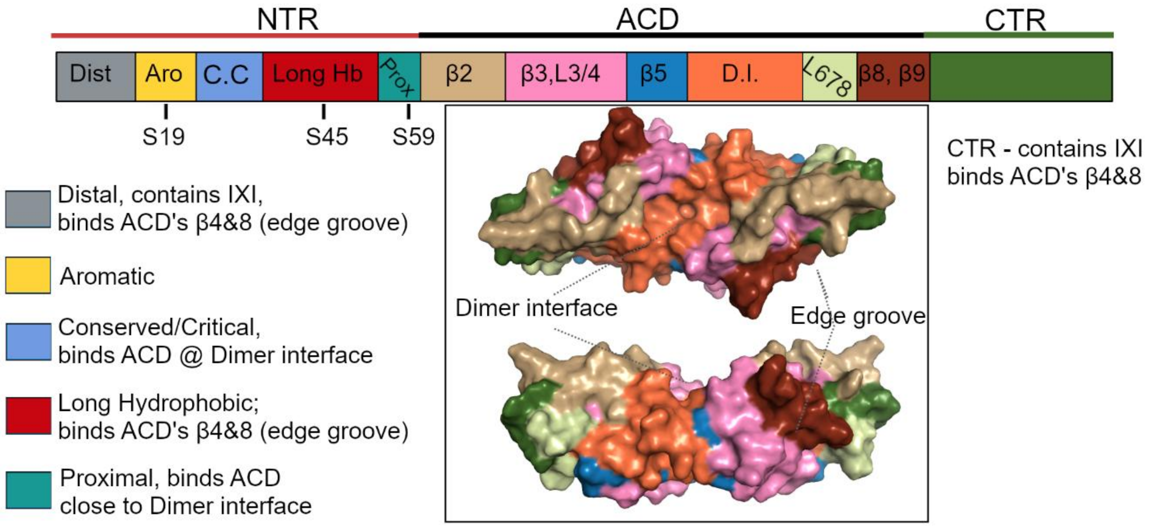


Figure S3. Schematic representation of the subregions within HSPB5. NTR and CTR are not modeled as they are intrinsically disordered. ACD subregions are denoted by secondary structure feature, β-sheet or loop, numbering. ACD dimer structure: PDB 2N0K.

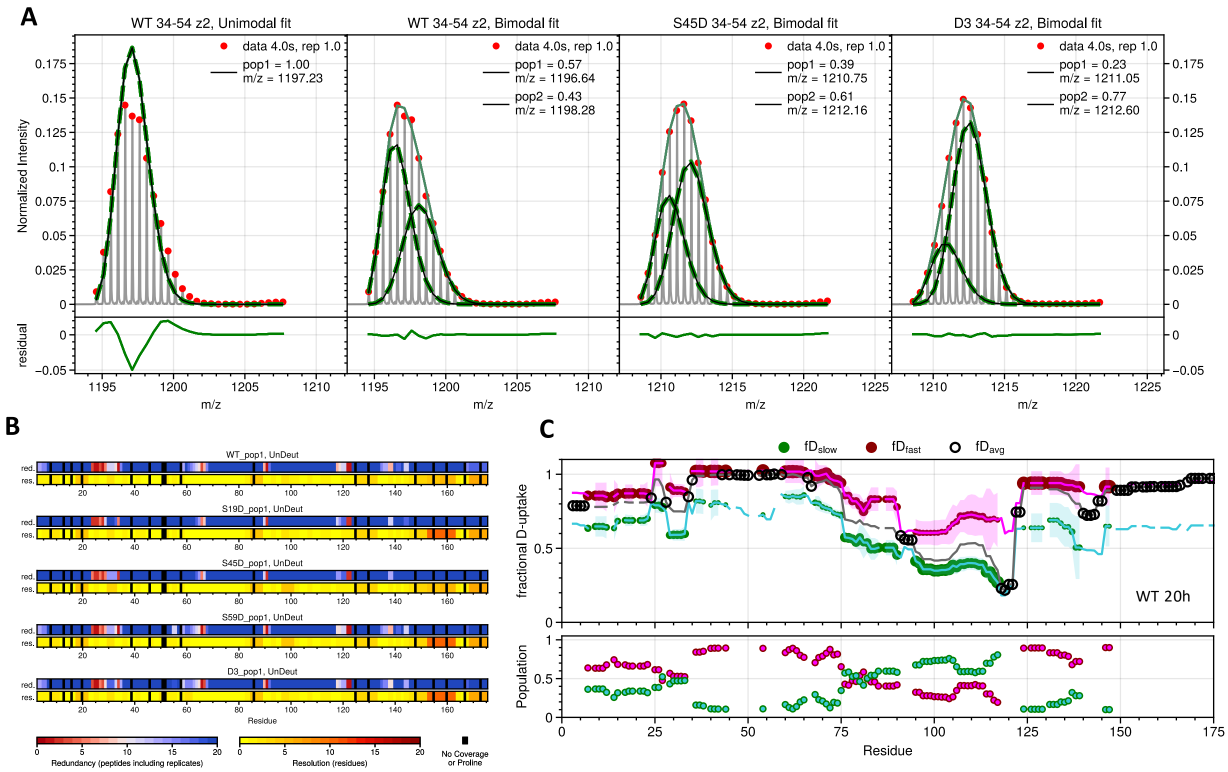


Figure S4. HDX-MS and bimodal behavior. A. HDX-MS spectra for peptide 34-54, charge 2+. The first panel shows that the unimodal fit for the WT data is poor resulting in a large residual (bottom). The second panel shows the same data using a bimodal fit, illustrating that the data is well described by a 2-state fit. This peptide is also well fit by 2-states for S45D and D3 (right panels). This peptide is in the Long Hydrophobic region of the NTR and exhibits different deuterium exchange behavior in WT, S45D, and D3. B. The redundancy and resolution plots for WT and each mutant. Red regions indicate areas that should be interpreted with care due to either a limited number of peptides covering that area (redundancy) or limited overlap of peptides in that region (resolution). C. Unimodal or bimodal behavior of WT-HSPB5 at 20h. Top, lines show the 1- or 2-state fits of the data; circles indicate whether the unimodal ($fD_{avg}$, black) or bimodal ($fD_{fast}$, magenta and $fD_{slow}$, green) fits are justified per residue. Bottom, corresponding populations ($p_{fast}$, magenta and $p_{slow}$, green) for residues with bimodal behavior.

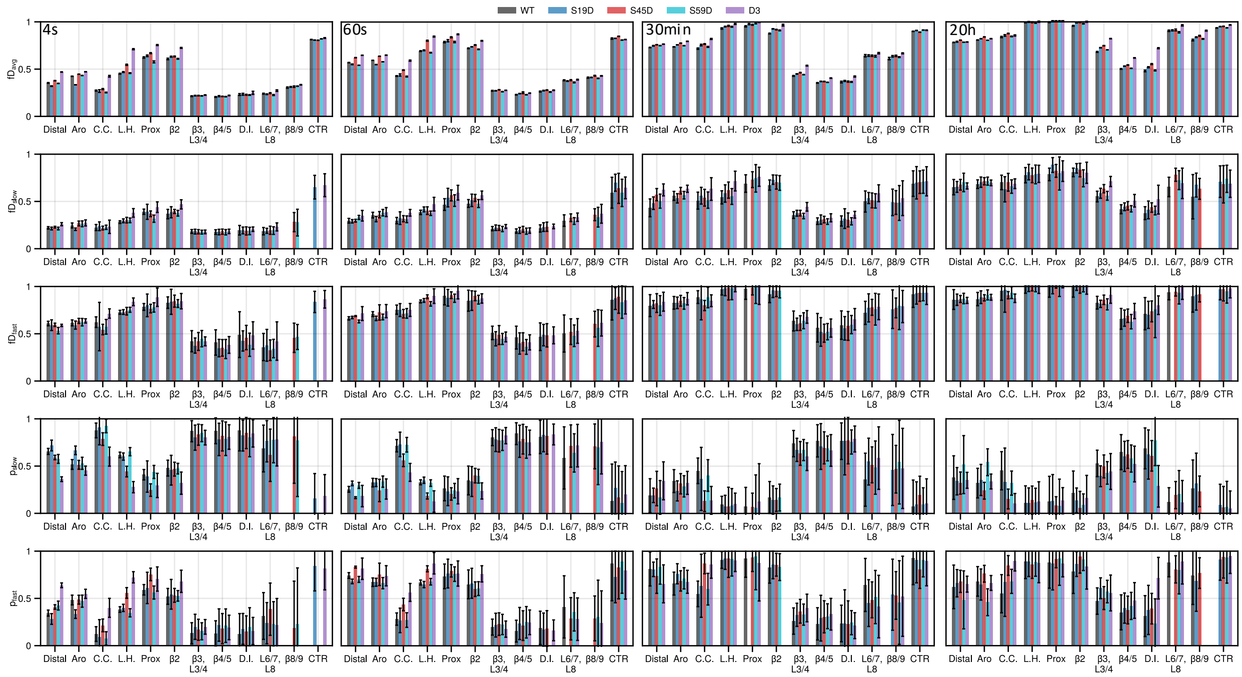


Figure S5. HDX-MS and bimodal behavior. Bar plots showing all parameters per region for all exposure times. Average values are for all residues, whereas bimodal parameters only include residues where that fit is justified.

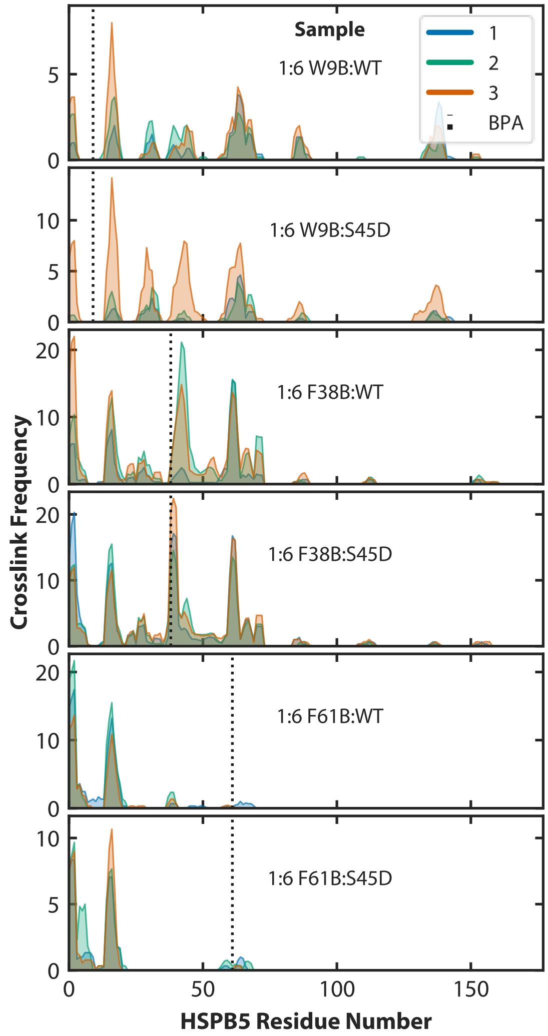


Figure S6. Counts of crosslinked spectra from individual replicates (blue, green, orange) identified between BPA variants W9B, F38B, and F61B of HSPB5 to WT or S45D HSPB5. BPA variants were individually diluted by WT or S45D in a 1:6 ratio. Crosslinks pictured proximal to BPA sites or between residues 35-56 for S45D are specific to the WT/S45D. Crosslinks were filtered to a 1% FDR. Crosslink counts for samples are as follows: W9B-WT: 58-95, W9B-S45D: 47-221, F38B-WT: 146-332, F38B-S45D: 298-321 PSMs, F61B-WT: 92-139, and F61B-S45D: 63-81 PSMs.
